# The probability of a unique gene occurrence at the tips of a phylogenetic tree in the absence of horizontal gene transfer (the last-one-out)

**DOI:** 10.1101/2024.01.14.575579

**Authors:** Nico Bremer, William F. Martin, Mike Steel

## Abstract

Gene loss is an important process in gene and genome evolution. If a gene is present at the root of a rooted binary phylogenetic tree and can be lost in one descendant lineage, it can be lost in other descendant lineages as well, and potentially can be lost in all of them, leading to extinction of the gene on the tree. In that case, just before the gene goes extinct in the rooted phylogeny, there will be one lineage that still retains the gene for some period of time, representing a ‘last-one-out’ distribution. If there are many (hundreds) of leaves in one clade of a phylogenetic tree, yet only one leaf possesses the gene, it will look like the result of a recent gene acquisition, even though the distribution at the tips was generated by loss. Here we derive the probability of observing last-one-out distributions under a Markovian loss model and a given gene loss rate *µ*. We find that the probability of observing such cases can be calculated mathematically, and can be surprisingly high, depending upon the tree and the rate of gene loss. Examples from real data show that gene loss can readily account for the observed frequency of last-one-out gene distribution patterns that might otherwise be attributed to lateral gene transfer.

## Introduction

Gene loss is an important and ubiquitous mechanism of genome evolution. In prokaryotes, gene loss acting on the whole genome is traditionally called reductive evolution (Andersson and Kurland, 1998; van Ham et al., 2003; Oshima et al., 2004; Hosokawa et al., 2006) and can result in miniscule genome sizes in parasites and endosymbiotic bacteria, the current record being *Macrosteles quadrilineatus* (Moran and Bennett, 2014) an endosymbiotic bacterium of leafhoppers that harbours only 137 protein-coding genes. Reductive evolution is also observed in symbiotic archaea (Waters et al., 2003) and in eukaryotes, especially among intracellular parasites (Tovar et al., 2003; Nicholson et al., 2022). Genome reduction through gene loss is also the *leitmotif* of genomic evolution in the endosymbiotic organelles of eukaryotic cells (Moore and Archibald, 2009), though many genes lost from organelle genomes have been transferred to the nucleus (Martin et al., 1998; Timmis et al., 2004). In eukaryotes, gene loss is also very common and widespread after whole-genome duplications (Blanc and Wolfe, 2004; Kellis et al., 2004; Brunet et al., 2006; Scannel et al., 2006). In general, if a gene belonging to a clade can be lost once in one lineage during evolution, it can be lost again in other lineages as well.

In comparative analyses, gene loss is easy to detect if losses are rare (Figure 1). If most genomes in a sample contain the gene, but one or a few do not, there can be little doubt that gene loss has occurred in the genomes lacking the gene. The more common loss is, the more difficult it becomes to distinguish from lateral gene transfer (LGT). If a given gene is present in about half of the genomes in a sample, the decision between loss and LGT becomes a matter of weighing the relative probabilities of LGT and gene loss, entailing an a priori assumption that LGT is roughly as common as loss. Many analytical tools to study prokaryotic genomes are currently in use that employ different and usually predetermined gain/loss ratios that are designed to differentiate between loss and LGT (Goodman et al., 1979; Page, 1994; Bansal et al., 2012; Szöllosi et al., 2013). In many cases, the overall average ratio of gene loss to LGT ends up being close to 1 in such applications, for obvious reasons. If loss predominates, then genomes steadily decrease in size across the reference tree (that is, ancestral genomes inflate), and if LGT predominates, genomes steadily increase in size across the reference tree (that is, ancestral genomes become too small) (Dagan and Martin, 2007). Some tools for estimating loss vs. LGT in current use can entail differences in loss vs. transfer probabilities for individual genes that differ by 20 orders of magnitude (Bremer et al., 2022).

**Fig. 1.**
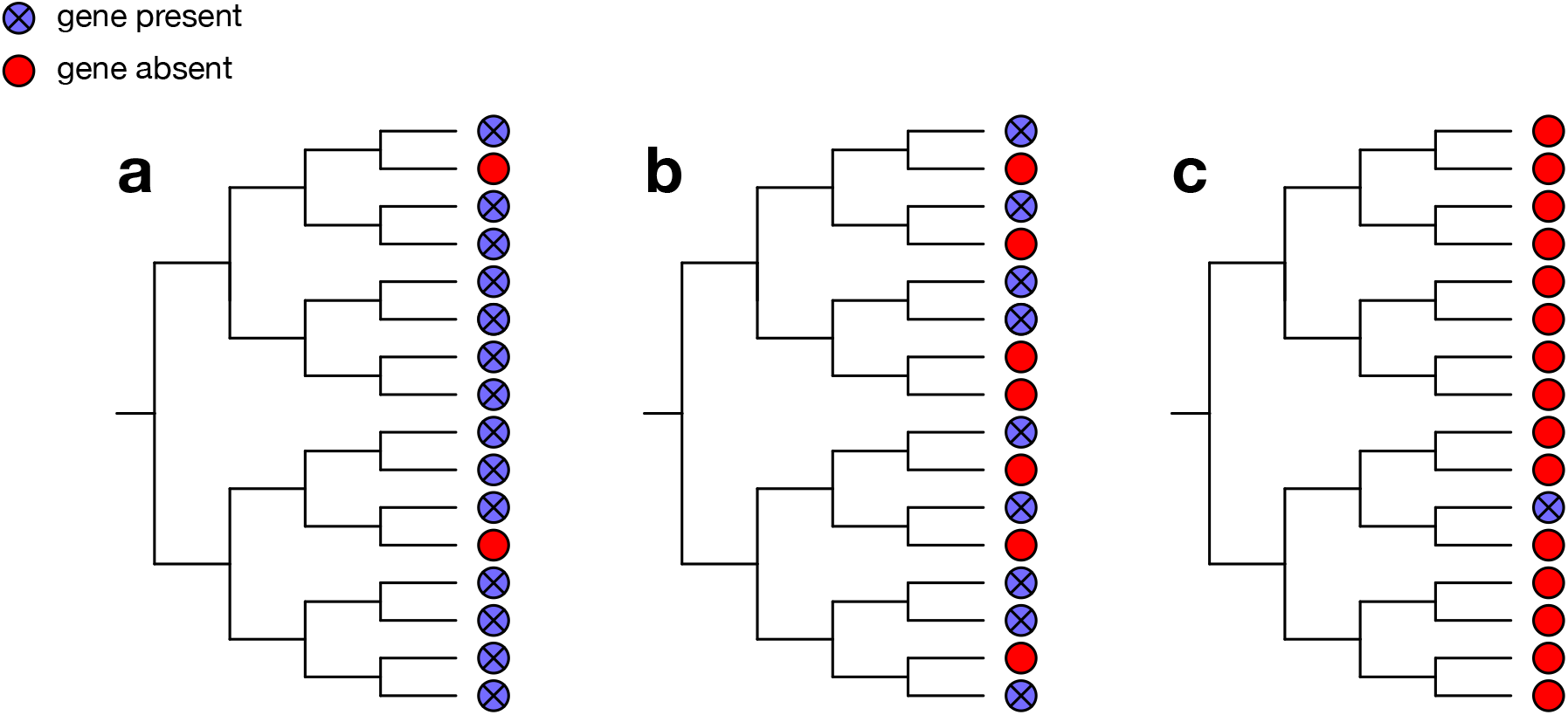
Hypothetical phylogenetic species trees showing the presence and absence of genes across all species in the trees. A blue circle with a cross indicates that the gene is present in this species, a red circle indicates that a gene is absent. (**a**) A distribution where gene loss most likely appeared on the branches to two species. (**b**) A case where the distribution of the genes that are present and absent is almost equal across the species tree. The decision between lateral gene transfer (LGT) and gene loss is highly dependent on the weighing of their relative probabilities. (**c**): illustrates a case where the gene is only present in one species. An easy (but not necessarily true) explanation for this would be LGT. This gene distribution across the tree can also be the result of a minimum of four gene losses if the gene was already present at the root node.

If gene loss is the predominant mode of genome evolution for a given gene in a given group, it will become lost in many lineages, ultimately in all. Just before the gene goes extinct in the group, however, there will exist a state in which the gene is present in only a few genomes, and finally, over time, only in one genome of the group. If this gene is in a eukaryote, but has homologs in prokaryotes, gene loss will produce a pattern that looks exactly like LGT: The gene is present in prokaryotes and one (or a few) eukaryotes. Under a loss-only mode of evolution, the last-one-out looks like an LGT, but the pattern was generated solely through gene loss. Here, we address the question of how likely it is to observe a last-one-out gene distribution under loss-only models.

## Mathematical modelling and algorithms

We now describe mathematical and computational methods to investigate the probability of last-one-out scenarios in both synthetic and real trees. We assume that each gene in a phylogeny can be lost along each lineage of a tree according to a continuous-time Markov process with loss rate *µ*, and which operates independently across genes and lineages.

### Recursions for a given tree

Let *T* be a rooted tree with a stem edge of length ℓ, and let *T*_1_, *T*_2_, … *T*_*k*_ denote the subtrees of *T* incident with this stem edge, as shown in Figure 2. Although the lengths of edges may correspond to time, and so be ultrametric, the algorithm described in this first section does not assume that edge lengths are ultrametric. Let 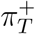 denote the probability that a gene *g* that is present at start of the stem edge of *T* is present in *exactly one* leaf of *T*, and let 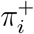 denote 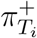 (the corresponding probabilities for the subtrees *T*_1_, …, *T*_*k*_). To calculate 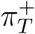 recursively, we also need to calculate the probability *π*_*T*_ that *g* is not present at any of the leaves of *T*, and we let *π*_*i*_ denote 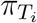 .

**Fig. 2.**
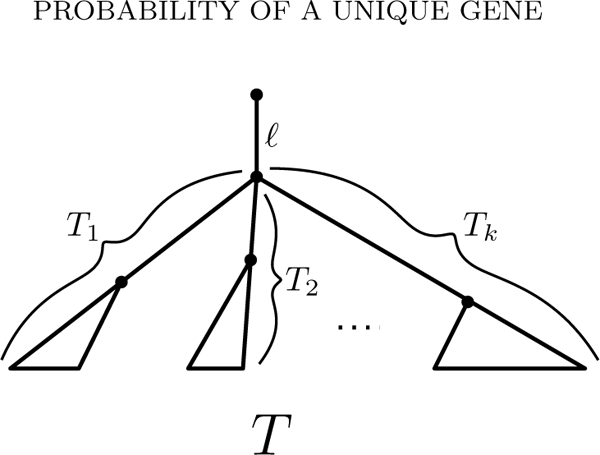

Note that if *T* consists of just a single stem edge of length *ℓ* (the base case in the recursion), then 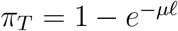 and 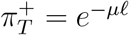. Thus we may suppose that *k* ⩾ 2. The following result (proved in the Appendix) provides a polynomial-time way to compute these quantities recursively via dynamic programming (progressing from the leaves to the root). Note that both Part (i) and (ii) are required for computing 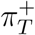.

Proposition 1 For the tree shown in Figure 2, the following recursions hold:

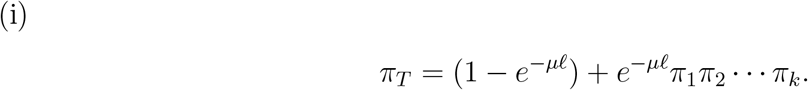

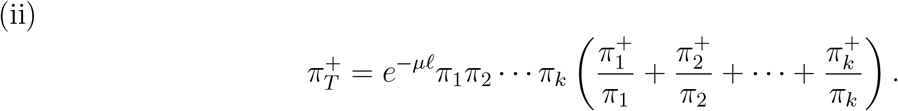

For binary trees, the second equation simplifies to:

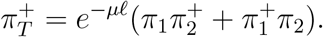

(iii) If there are *G* ⩾ 1 genes present at the top of the stem edge of *T*, the number of genes that appear in just one leaf of *T* has a binomial distribution with parameters 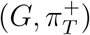.

To illustrate Proposition 2 with a simple example, consider the tree in Figure 2, where each of the subtrees *T*_1_ …, *T*_*k*_ is a single leaf at the same distance from the root, and ℓ = 0 (the ‘star tree’). Under the gene-loss model, a gene that is present at the root of the tree will be present at exactly one leaf of this tree precisely if there are *exactly k* − 1 loss events. This might seem very unlikely for large values of *k*. However, if *µ* is chosen carefully, then the probability of this event can be at least *e*^*−*1^ = 0.367 regardless of how large *k* is. Nevertheless, if we consider the posterior value of this probability by taking a uniform prior on 1 *−e*^*−µ*^ (setting the height of the tree to 1), then this posterior probability tends to 0 as the number of leaves of the tree (*k*) grows. The proof of these claims and the analysis of this star tree when we allow ℓ *>* 0 are provided in the Appendix. Of course, the star tree is a highly non-binary tree, which raises the question of whether 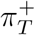can be close to *e*^*−*1^ when *T* is binary and the number of leaves is large. This is indeed possible: we can simply resolve the polytomy at the root by using very short interior edges to obtain a binary tree for which 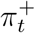 will be close to the corresponding value for a star tree and hence can be close to *e*^*−*1^ for a suitably chosen value of *µ*. However, for trees generated by simple phylodynamic models, this is no longer the case, as we demonstrate in the next section.

### Random trees

Suppose now that *T* is generated by a standard birth–death model (Kendall, 1948; Lambert and Stadler, 2013) with speciation rate *λ* and extinction rate ***ν***, starting from a single lineage at time *t* in the past. The tree *T* is now a random variable, denoted *𝒯*_*t*_, and the number of species at the present (denoted *N*_*t*_) is also a random variable and has a (modified) geometric distribution with expected value 𝔼[*N*_*t*_] = *e*^(*λ−****ν***)*t*^. We will suppose that *λ >* ***ν*** since otherwise the tree *𝒯*_*t*_ is guaranteed to die out as *t* grows. Let 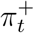 be the probability that a gene *g* that is present at start of the stem edge of *𝒯*_*t*_ is present in *exactly one* leaf of *𝒯*_*t*_. The following result precisely describes the maximum value that 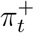 can take as *µ* (the rate of gene loss) varies over all possible positive values. The short proof is provided in the Appendix.

Proposition 2

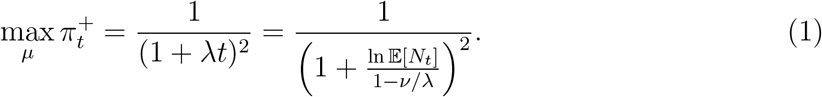

Notice that although 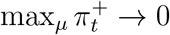for Yule trees as they grow in their expected size, the convergence is quite slow as a function of the expected number of leaves of the tree, due to the presence of the logarithmic function on the right of Eqn. (1). Also, if there are *G* ⩾ 1 genes present at time 0, then the expected number of genes that will be present in just leaf of *𝒯*_*t*_ is 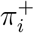. However, in contrast to Proposition 1(iii), the number of genes present in just one leaf of *𝒯*_*t*_ is no longer binomially distributed, since this number is now a compound random variable because it is dependent on the random variable *𝒯*_*t*_.

To illustrate Proposition 2, consider (pure–birth) Yule trees (i.e., ***ν*** = 0) with an expected number of 150 leaves. Then max_*μ*_ 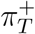≈0.028, and so for 10,000 independent genes and this optimal rate of gene loss, the expected number of genes that would be last-one-out (i.e., present in just one leaf of these Yule trees) would be around 280. This provides some insight into the results described in the next section.

## Analysis of real genome data

To test this algorithm on real genome data we chose the example of genes in eukaryotic genomes that have homologues in prokaryotes but that are present in only one or a few eukaryotic lineages. Such patterns are taken as evidence for the workings of differential loss, under the assumption that loss will generate such patterns (Ku et al.2015), or as evidence for the workings of LGT (Cote-L’Heureux et al., 2022) under the assumption that LGT rather than loss generates such patterns. The calculation of the probability of a gene being present at the root node and remaining in exactly one leaf of a eukaryotic tree requires a rooted species tree and a gene loss rate *µ*. Reconstructing a eukaryotic species tree is challenging, and there is currently no consensus on the position of the root (Keeling and Burki, 2019; Burki et al., 2020). Although the loss rates can be adjusted and averaged across a range of values, the backbone trees with all their nodes, branches and branch lengths are not that easily adjustable.

We therefore analyzed a set of ten eukaryotic gene trees with 150 leaves each. These gene trees need not be representative of the true phylogeny of eukaryotes, nor need they show a pattern of gene distribution that could be indicated as LGT. The different trees were selected merely to show that different phylogenies can have an influence on the calculated probability of a gene being present at the root node and remaining only in one leaf of a eukaryotic tree. Furthermore, the different gene trees with 150 leaves provide an opportunity to estimate the overall probability of observing a last-one-out pattern if we consider thousands of eukaryotic genes with prokaryotic homologs (Figure 3). In Figure 3, we assume that 10,000 genes were present in the last common ancestor of 150 eukaryotes. For those trees, the mean probabilities result in 232 (lowest mean) to 790 (highest mean) last-one-out cases that look like LGT but actually are the result of differential loss in a loss-only mode of evolution for a 10,000 gene ancestral genome. Looking at the median, we would find 11 (lowest median) to 755 (highest median) cases, depending on the tree itself. Since loss rates are not constant over time, we cannot assume that these percentages resemble the ‘real’ amount of those cases due to differential loss. What they do show, however, is that last-one-out cases are not so rare that they can be excluded a priori. If the loss rate is ideal, meaning that the maximum probability of last-one-out cases for the given tree is achieved, we would see between 532 (lowest maximum) and 2,502 (highest maximum) out of the 10,000 genes resulting in a last-one-out scenario, which is a substantial frequency. That is, in a study of 10,000 gene families present in the eukaryotic common ancestor, one would expect to observe dozens, hundreds, or even thousands of last-one-out patterns in trees sampling 150 genomes obtained solely as the result of differential loss. These cases would appear, in a gene phylogeny, as a single eukaryote (or group thereof) branching within prokaryotic homologues.

**Fig. 3.**
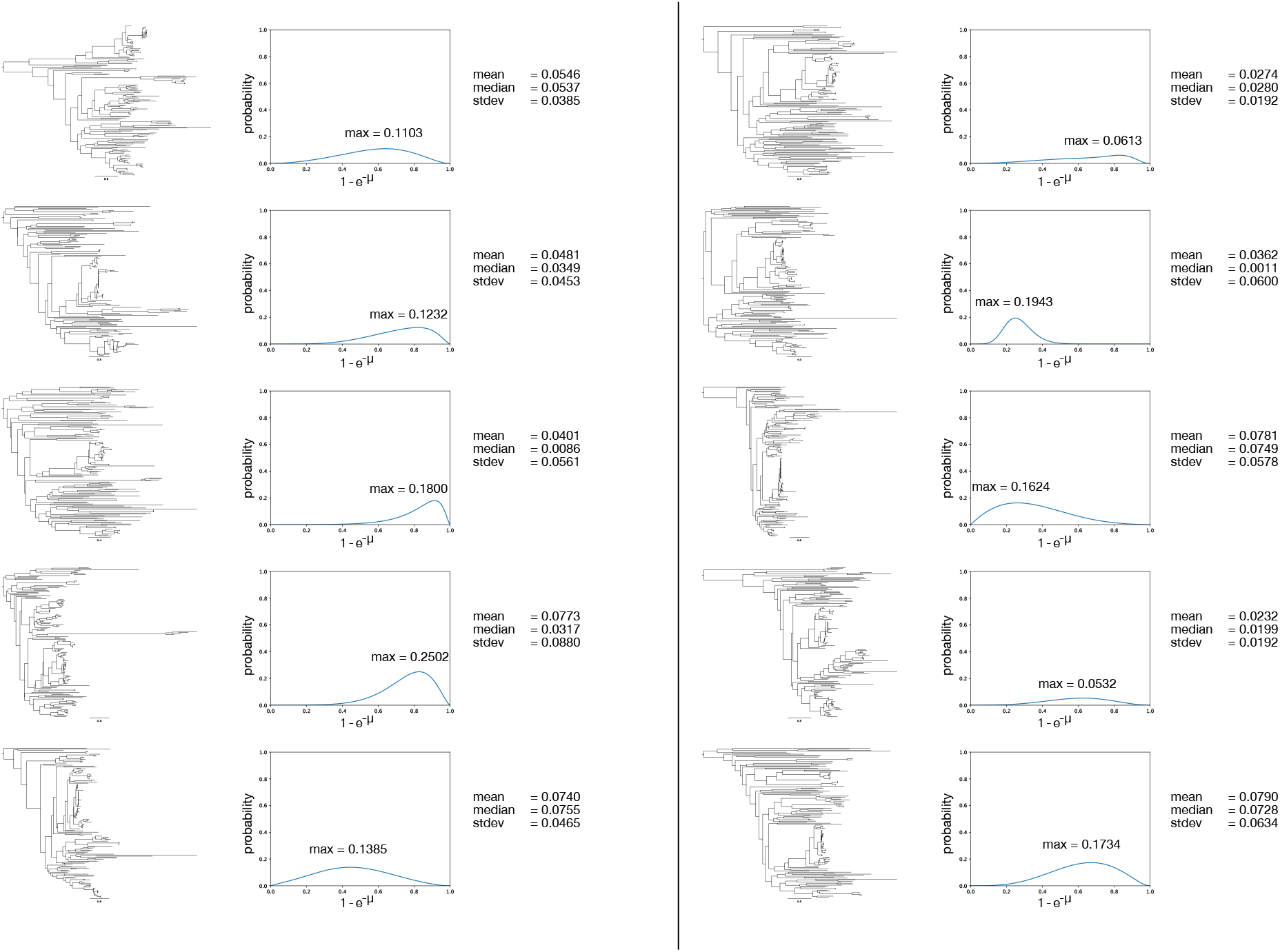
Ten eukaryotic gene tree phylogenies with 150 leaves each and the corresponding probabilities for a last-one-out scenario against 1 *− e*^*−µ*^ (*µ* = gene loss rate). The trees show various possibilities of species trees without assuming that those trees represent a real eukaryotic backbone tree. They show that the phylogeny itself has an influence on the probability of a last-one-out scenario, but that the overall probability is comparably high.

## Reinspecting some eukaryote LGT claims having last-one-out topologies

The surprisingly high probability to observe a gene that is present in the root node and only in one species or clade and lost in all other leaves of a tree offers a new approach to investigate data that looks like evidence for LGT based on a rare or sparse gene distribution. Differential loss can–and will–produce last-one-out patterns that look just like lineage specific LGT. It is therefore possible, if not probable, that many reports claiming evidence for LGT are in fact due to differential loss.

A recent study provides a case in point. Cote-L’Heureux et al. (2022) looked for lineage-specific presence of prokaryotic genes in eukaryotes that would provide the strongest possible evidence, in their view, for the workings of LGT from prokaryotes to eukaryotes. They sampled 13,600 gene families, 189 eukaryotic genomes and 540 eukaryotic transcriptomes, looking for recent lineage-specific LGT (topologies that we call last-one-out patterns). Among the 13,600 eukaryotic gene families sampled, they found approximately 94 putative cases of LGT that represent a last-one-out pattern, that is, a restricted single-tip distribution of a prokaryotic gene in a eukaryotic genome or group, which they interpreted as strong evidence for LGT. Our present findings (Figure 3) indicate that in Cote-L’Heureux et al. (2022) the number of cases identified in their study (94) is very close to the lower bound of the expectations for last-one-out topologies of similarly sized data sets, in which all the last-one-out topologies can be accounted for by differential loss alone, with no need to invoke LGT.

One clear prediction of lineage-specific LGT versus loss for last-one-out cases is this: If lineage-specific acquisition is the mechanism behind the observed rare presence pattern for a eukaryotic gene, then the acquisition would need to be evolutionarily late (i.e., a tip acquisition). That is, the prokaryotic donor and the eukaryotic gene should share a higher degree of sequence similarity, on average, in comparison to genes that trace back to the eukaryotic common ancestor. This is the reasoning behind the analysis of Ku et al. (2015) and Ku and Martin (2016), who looked for evidence of recent acquisitions of prokaryotic genes in sequenced eukaryotic genomes. Ku et al. (2015) found that, in eukaryotic genomes, rare genes that have prokaryotic homologs were not more recently acquired (more similar to prokaryotic homologs) than genes that trace back to the eukaryotic common ancestor, suggesting that their rare occurrence is the result of differential loss rather than lineage-specific acquisition (Ku et al., 2015; Ku and Martin, 2016) (Figure 4a,b).

**Fig. 4.**
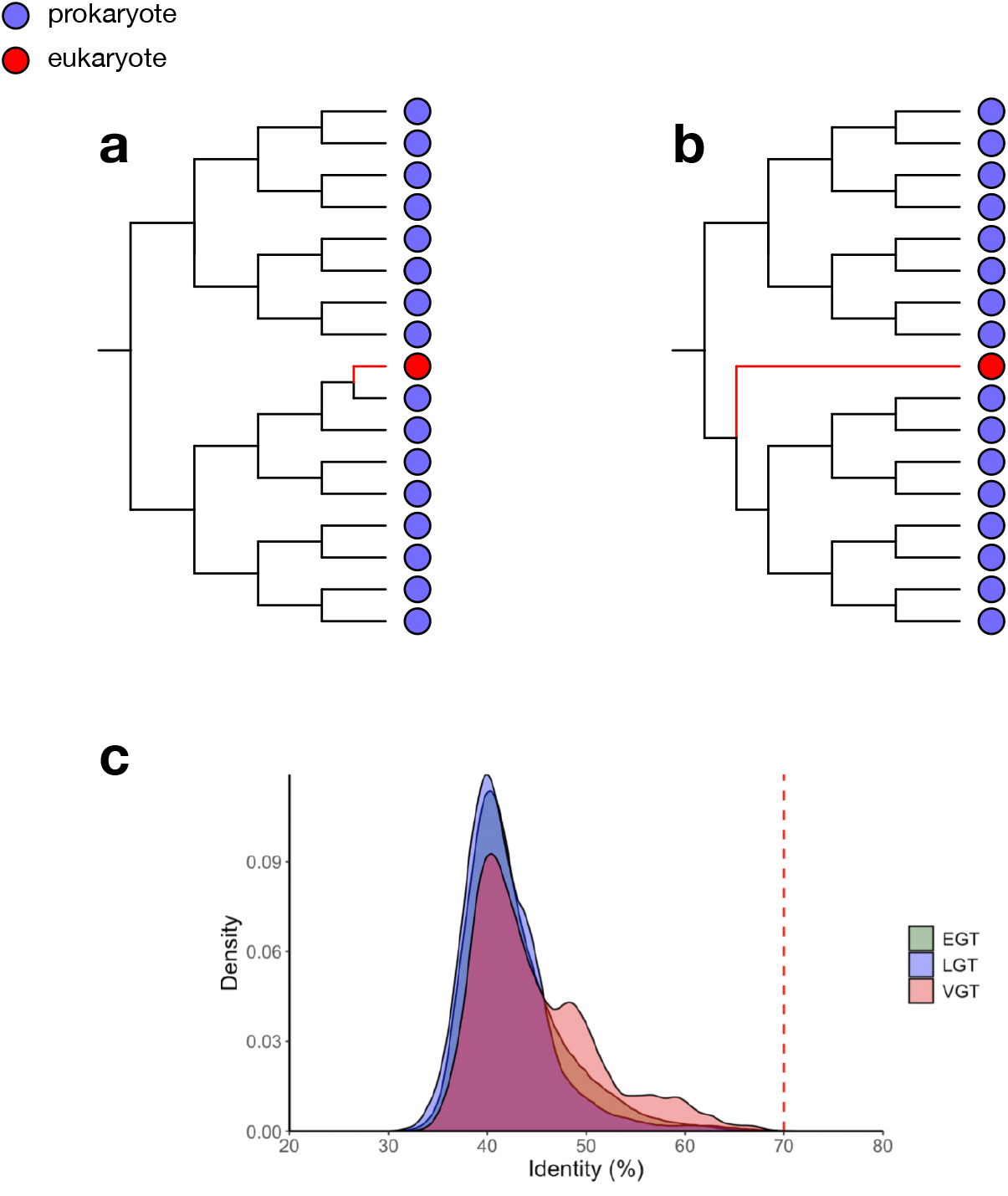
Similarity of eukaryotic last-one-out cases to prokaryotic homologs. (**a**) Phylogenetic distribution of genes where the eukaryotic gene is considered to be the result of LGT due to its high similarity to one prokaryotic homolog. (**b**) The eukaryotic gene does not have a substantially high similarity to its prokaryotic homologs. It can therefore not be the result of recent LGT and is more likely the result of differential gene loss. (**c**) Supplementary Figure 9 from Cote-L’Heureux et al. (2022) showing that genes assumed to be the result of LGT are at most 70% similar to their prokaryotic homologs. This finding supports the ‘70% rule’ of Ku and Martin (2016) and furthermore shows that these cases are more likely to be the result of differential loss instead of LGT. EGT: endosymbiotic gene transfer (genes acquired from chloroplasts or mitochondria); LGT: lateral gene transfer; VGT: vertical gene transmission.

Cote-L’Heureux et al. (2022) employed the same test, making the same kind of comparison that Ku et al. performed, namely, they looked for cases in which the prokaryotic gene was acquired recently by the eukaryotic lineage, using the criterion of sequence similarity. What they found was the distribution shown in Figure 4c, namely that the cases they suspected to be LGTs were just as old, in terms of sequence divergence, as genes that were acquired from the mitochondrion. In other words, there were no obviously recent acquisitions, as all of the prokaryotic genes that they interpreted as recent LGTs had the hallmark of ancient acquisition, just as Ku et al. (2015) suggested. Cote-L’Heureux et al. (2022) offered no explanation for the finding that genes they interpreted as recent acquisitions via LGT were just as ancient, in terms of sequence identity, as genes acquired from mitochondria (Fig. 4c). One interpretation is that the genes in their LGT class were not LGTs after all but were the result of differential loss instead. Differential loss directly explains why such genes show just as much sequence divergence to prokaryotic homologues (Ku et al. 2015) (Figure 4c) as genes present in the eukaryotic common ancestor. LGT models would need to invoke an ad hoc corollary assumption of substitution rate acceleration for every gene with a last-one-out pattern to account for the absence (Figure 4c) of eukaryotic LGTs having high (*>* 70%) sequence similarity to prokaryotic homologs. Differential loss requires no rate acceleration corollary. Furthermore, the model presented here closely predicts the frequency of observing last-one-out patterns under a variety of topologies and loss rates.

## Conclusion

Sparse gene distributions in eukaryotes are often interpreted as evidence for gene acquisition via LGT from prokaryotes. However, gene loss can generate the same patterns, and estimates for the probability of observing a single gene at the tip of a phylogenetic tree as the result of differential loss within a given clade, as opposed to LGT, have been lacking. Here, we have derived the probability of observing such cases, which we call last-one-out patterns, because under a loss-only model, the last gene to be lost looks like an instance of LGT. The probability depends on the size and shape of the tree, and the loss rate *µ*. We find that the probability of observing a last-one-out topology can be (surprisingly) high. A simple algorithm applied to simulated eukaryotic trees provides estimates for the frequency of last-one-out patterns resulting from a loss only model that are slightly higher than, but generally in good agreement with, observations from a recent study in which all last-one-out topologies were interpreted as evidence for LGT. Gene loss is a prevalent process in genome evolution. If one lineage can lose a given gene, others can as well. Gene loss can, and does, generate patterns that look just like LGT. Even for large data sets, the probability of last-one-out topologies can be surprisingly large, because, depending upon the tree, the number of losses required to account for a last-one-out topology can be small.

## Acknowledgements

This project has received funding from the European Research Council (ERC) under the European Union’s Horizon 2020 research and innovation programme (grant agreement No. 101018894). MS thanks the Alexander von Humboldt Foundation for supporting his visit to Germany in 2023.

## Data availability

Phylogenetic gene trees are available as Supplemental Data under https://doi.org/10.6084/m9.figshare.24980901.v1.

## Appendix: Mathematical details

### Proof of Proposition 1

Both parts involve straightforward applications of the law of total probability and conditional independence (due to the Markovian nature of the model).

Part (i): A gene present at the top of the stem edge but at none of the leaves has either disappeared on the stem edge of length ℓ, (an event that has probability 1 *− e*^*−µℓ*^) or it is present at the end of the stem edge (with probability *e*^*−µℓ*^) and is not present in any leaf of the *k* subtrees, *T*_1_, … *T*_*k*_. Since these *k* latter events are independent, we can multiply their probabilities to obtain the required joint probability.

Part (ii) follows by considering the *k* ways in which a gene present at the top of the stem edge can be present in exactly one of the leaves of *T* (depending on which trees *T*_1_, … *T*_*k*_ that this leaf appears in). The term *e*^*−µℓ*^ ensures that the gene survives to the other end of the stem edge, and each of the *k* summation terms involves the gene being present in exactly one leaf of exactly one of the subtrees (say *T*_*i*_) with probability 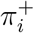 and not present in any leaf of the other subtrees (with probability *π*_*i*_ for *i /*= *k*). Again, by independence, we can multiply these probabilities together.

Part (iii) follows from the assumptions that gene loss events are independent and that the tree on which they take place is fixed.

### Analysis of the star tree

Consider the star tree *T* with *n* leaves, with no stem edge (i.e., ℓ, = 0), and let *y* = 1 *− e*^*−µt*^. In this case, by Proposition 1, we have: 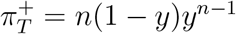. Solving the equation 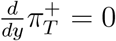 gives *y* = 1 *−* 1*/n*, and for this value of *y* we obtain the maximal value of 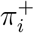, namely (1 *−* 1*/n*)^*n−*1^. For example, for *n* = 4 this gives *y* = 3*/*4 and 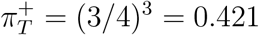 . Notice that as *n* grows, 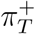 converges to *e*^*−*1^ = 0.367…

Now suppose we set *t* = 1 and consider a uniform prior on 1 *−e*^*−µ*^. Let *Y* denote the uniform random variable on [0, 1]. Then *Y* = 1 *− e*^*µ*^ and so *µ* = *−* ln(1 *− Y*). It follows that the the density function for *µ* (denoted *f*_*µ*_) is given by *f*_*µ*_(*x*) = *e*^*−x*^ for *x >* 0. Conditional on *µ* = *x* we have 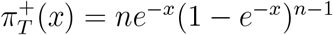, and so the expected value of 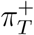 is:

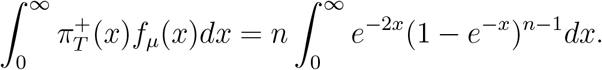

If we now set *u* = 1 *− e*^*−x*^, this expression becomes 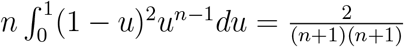 which tends to 0 at an inverse quadratic rate as *n → ∞*.

Next, consider the slightly more general setting of a tree *T* that has an edge from its root to a star tree with *n* leaves, and with an additional leaf adjacent to the root (thus this tree also has *n* + 1 leaves in total). Without loss of generality, let the edges of the star tree each have length 1, and let the stem edge connecting it to the root of *T* have length *𝓁*. Thus, the tree has height *𝓁* + 1. We have: 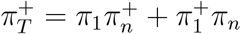, where *π*_*n*_ refers to the startree, and *π*_1_ refers to the leaf incident with the root. We have: *π*_1_ = 1 *− e*^*−µ*(1+*ℓ*)^ and 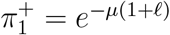Furthermore,

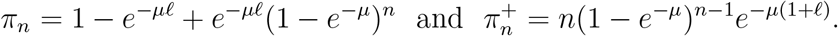

Thus,

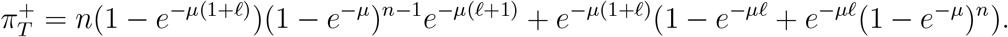

For example, when *n* = 2 and ℓ = 2, 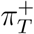has a maximal value of 0.326 (as *µ* varies). For *n* = 3 and *𝓁* = 5, 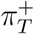 has a maximal value of 0.231.

### Proof of Proposition 2

Consider a birth–death tree with speciation and extinction rates *λ* and *µ*, respectively (with *λ > µ*), grown for time *t* from a single individual at time 0. On this tree, superimpose a continuous-time Markov process of gene loss along the branches of the tree, starting with an initial single gene present at time 0. Let *X*_*t*_ (*t ⩾* 0) denote the number of leaves of the tree (at time *t*) that are carrying the initial gene. Then *X*_*t*_ is described by a birth–death process with birth rate *λ* and death rate *θ* = *µ* + ***ν***. Consequently,

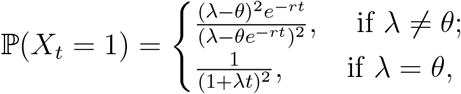

where *r* = *θ − λ* (Kendall, 1948). Now, 𝕡(*X*_*t*_= 1) = *φ*^2^, where 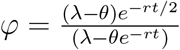 . Therefore, to find the maximal value of 𝕡(*X*_*t*_ = 1) = *φ*^2^ as we vary *µ ⩾* 0 (recalling that ***ν*** *< λ*), we solve the equation:

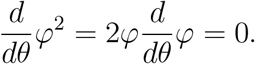

This leads to the solution *θ* = *λ*, which provides the unique value that maximises *φ*. Straightforward algebra then leads to the claimed result.

